# Circular 23S rRNA within archaeal ribosomes

**DOI:** 10.1101/2025.04.27.650855

**Authors:** Ling-Dong Shi, Petar I. Penev, Amos J. Nissley, Dipti D. Nayak, Rohan Sachdeva, Jamie H. D. Cate, Jillian F. Banfield

## Abstract

The ribosomal RNAs (rRNAs) that form the core of ribosomes are believed to occur as linear molecules. Here, we investigated rRNAs from diverse and mostly uncultivated archaea and found evidence that, in at least eight phylum-level archaeal lineages, the 23S rRNAs within mature ribosomes are circular. Sequencing of total cellular RNA indicates that the transcriptional abundances of archaeal circular 23S rRNAs vastly exceed those of linear counterparts, and linear versions are often undetectable. As the majority of rRNAs derive from mature ribosomes, the data suggested that the circular molecules are within the ribosomes. Thus, we directly sequenced RNA extracted from isolated ribosomes of a model archaeon, *Methanosarcina acetivorans*, and confirmed that the assembled 23S rRNAs are circular. Structural modeling places the 5’ and 3’ ends of the linear precursors of archaeal 23S rRNAs in close proximity to form a GNRA tetraloop (in which N is A, C, G, or U and R is A or G), consistent with their further circularization within ribosomes. We confirm the existence of circular 16S rRNA intermediates in transcriptomes of most archaea, yet a circular form is not evident in some distinct archaeal groups. Overall, the results uncover unexpected variations in the processing required to generate mature rRNAs and the conformation of functional molecules in archaeal ribosomes.

## Introduction

Translation is less well studied in Archaea than in the other Domains of life, yet this process is of great interest because archaeal systems are more similar to those of Eukaryotes than those of Bacteria^1–3^. Once transcribed, ribosomal RNAs (rRNAs) must be processed to maturation before assembly into ribosomes^4–6^. Several studies of pure cultures demonstrate that archaeal rRNA processing often involves excision of individual rRNAs from polycistronic rRNA precursors and circularization of the excised rRNAs. It is widely believed that re-linearization of circular rRNA intermediates occurs prior to the incorporation of rRNAs into the ribosome^7–12^. However, all the studied isolates are confined to Euryarchaeota and Crenarchaeota, leaving open the possibility that different processes occur in other archaeal groups.

Since the advent of genome-resolved metagenomics^13^, culture-independent *in silico* analyses have greatly increased genomic sampling of Archaea and brought to light innumerable lineages that were not known based on laboratory cultures^14–17^. More recently, environmental microbiomes have been studied by directly sequencing extracted RNA, an approach referred to as metatranscriptomics^18^. The messenger RNA (mRNA) sequences are used to document *in situ* microbial activity, typically after depletion of the rRNAs that comprise the vast majority of transcriptomes and hinder the resolution of the mRNA pool^19^. However, the rRNAs can be used to investigate the steps by which rRNAs are processed in diverse, usually uncultivated organisms and to predict the form of mature molecules within ribosomes.

Here we performed short- and long-read deep DNA and RNA sequencing to access the genomes and transcriptomes of diverse and mostly uncultivated archaea in complex wetland soil samples. We mapped the rRNA transcripts to genomes and used read alignment discrepancies to document circularization. Results of total cellular RNA sequencing indicated that 23S rRNAs from eight phylum-level archaeal lineages are mostly in circular forms, suggesting that ribosome-associated 23S rRNAs are circularized. We augmented these analyses with direct sequencing of rRNAs from ribosomes isolated from a model archaeon, *Methanosarcina acetivorans*, and concluded that the investigated archaea have circular 23S rRNAs within their ribosomes. We also uncovered evidence suggesting that the steps in processing 16S rRNAs vary across archaeal lineages. These findings expand our understanding of rRNA processing and the structure of ribosomes in the Archaeal Domain of life.

## Results and Discussion

DNA and RNA were extracted from samples of wetland soil collected from a site in Lake County, California, USA. Genome-resolved metagenomic data have revealed abundant, diverse, and mostly uncultivated archaea in this soil ecosystem, including methane-oxidizing archaea, methanogenic archaea, Asgard archaea, and their associated extrachromosomal elements^20–23^. Here, we performed metatranscriptomic sequencing to determine the structures of archaeal rRNAs and provided insights into archaeal rRNA processing.

A circularized *Methanoperedens* genome was reconstructed using long-read PacBio data and completion was confirmed using short Illumina reads. Based on the genome sequence, we predicted the locations of the rRNA genes. DNA reads spanned the transition from intergenic regions into the rRNA genes (Supplementary Fig. 1a). We then mapped Nanopore and Illumina transcript reads to the *Methanoperedens* genome and observed that reads placed at the starts and ends of 16S and 23S rRNA genes only partially agreed with the reference sequence. The discrepant portions of the transcript reads from the region preceding the genes exactly matched the ends of the genes and *vice versa*, indicating that these reads derived from circularized rRNAs (Supplementary Figs. 1b-1c). As this method does not require pure cultures and can document the various rRNA forms present (Fig. 1), we used it to investigate whether rRNA processing that involves a circularization step occurs in other archaea.

**Fig. 1.**
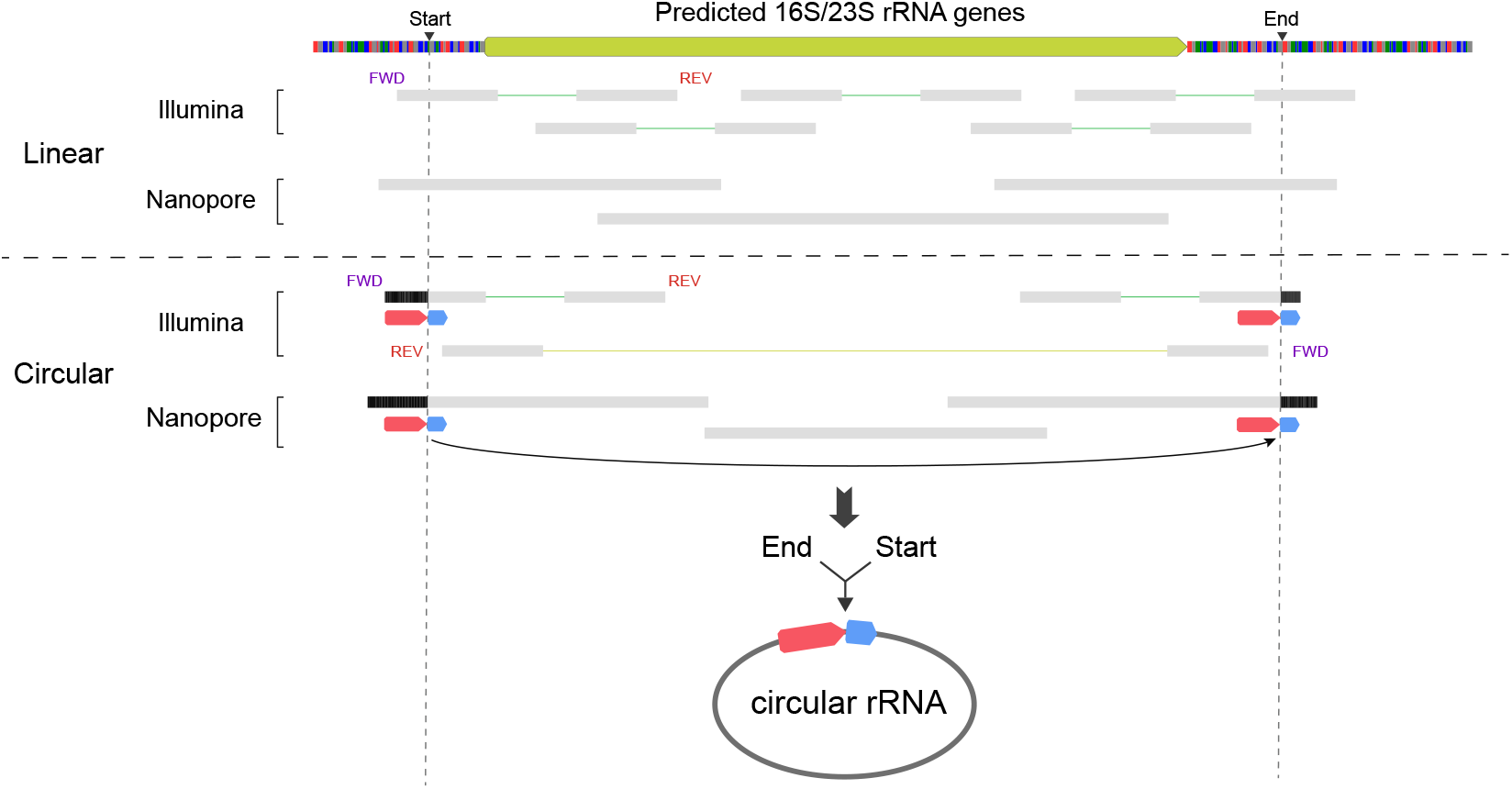
Diagram of transcript mapping to predict rRNA forms. Black triangles on the top genome reference indicate inferred start and end sites of circular rRNAs. Green lines denote the forward orientation of Illumina paired reads (i.e., FWD -> REV) while yellow lines indicate the reverse orientation of them (i.e., REV -> FWD). Gray bars are mapped reads. Colored arrows denote identical sequences. Black parts inside reads show mismatches to genome references.

We analyzed all 16S rRNA genes from the wetland soil dataset and assigned taxonomic classifications to each at the highest resolution possible. Long-read PacBio HiFi sequences from several samples were used to capture entire rRNA gene sequences (Supplementary Fig. 2). In total, 11,710 16S rRNA sequences ≥ 500 bp were assembled. The 4,580 full-length and potentially complete 16S rRNA genes were clustered into 2,576 species-level groups at 99% identity (Supplementary Fig. 3)^24^. Of the 1,125 16S rRNA genes with sequence coverage in start and end regions needed to evaluate circularization, 963 were from Bacteria. As expected based on prior studies of bacterial rRNA processing^4^, no evidence supported the presence of circularized bacterial 16S rRNA transcripts. The remaining 162 genes were from Archaea. Of these, 78 showed evidence that a fraction of the 16S rRNA transcripts were in circularized forms. Archaea with circular 16S rRNAs are phylogenetically distributed in eight phyla (Fig. 2), including lineages that currently do not have representative isolates (e.g., Bathyarchaeota), thus expanding the diversity of microorganisms that are capable of circularizing 16S rRNAs. Circularization was also observed for 23S rRNAs in all investigated archaea.

**Fig. 2.**
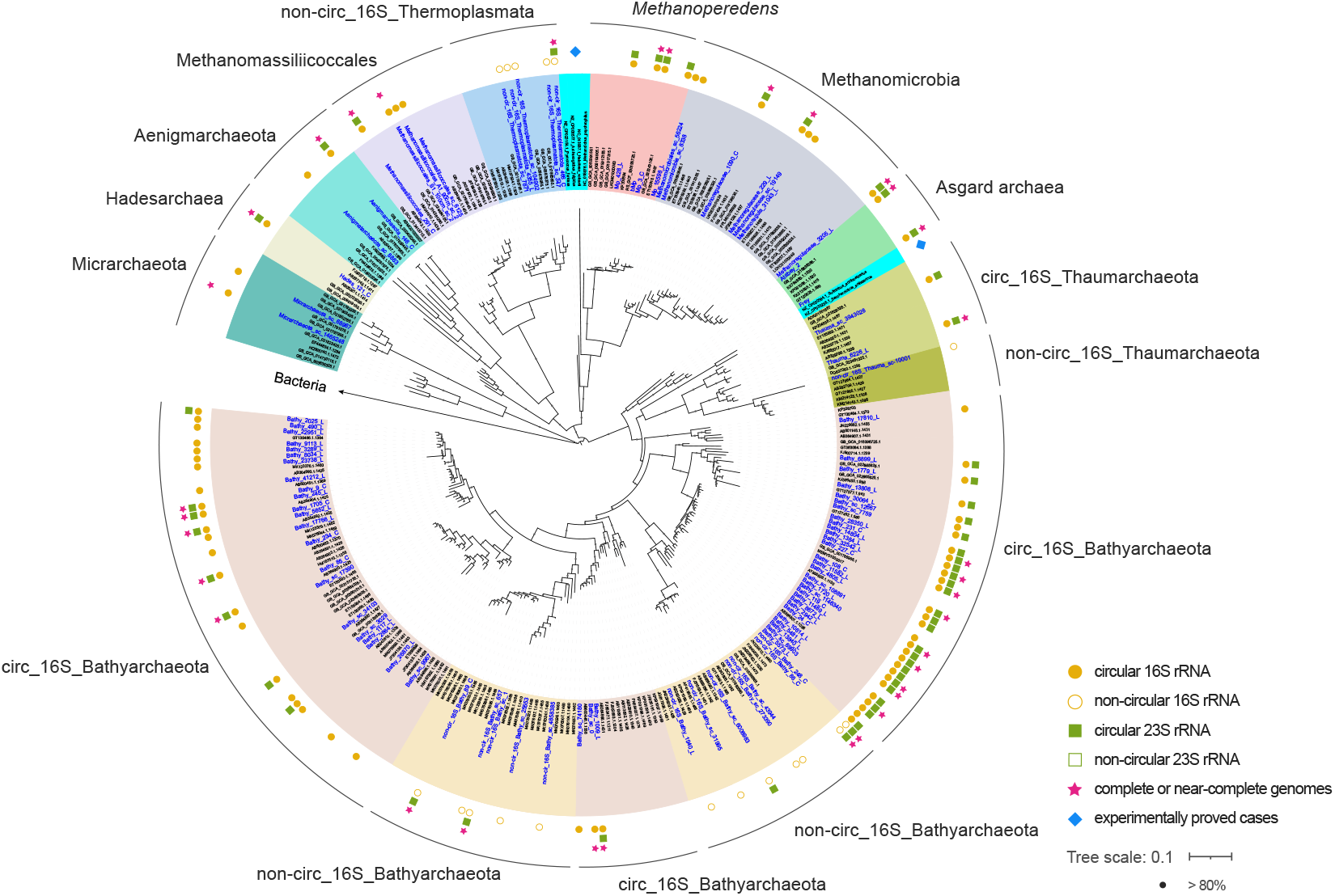
Phylogeny of archaeal 16S rRNA genes with confident determination in rRNA forms. Sequences highlighted in blue were identified in this study. The tree was rerooted using Bacteria (*Escherichia coli* and *Bacillus subtilis*) as the outgroup. Filled/open circles and rectangles beside genomes indicate observed circular/non-circular 16S rRNAs and 23S rRNAs, respectively. Stars represent potentially complete or near-complete genomes, the latter of which include most single-copy genes and < 20 scaffolds. Diamonds show cases that are experimentally proven to circularize 16S and 23S rRNAs. Support values were calculated based on 1000 replicates.

Archaeal 16S rRNA transcripts that circularize exhibit an extension of 105.6 ± 22.2 bp (n=78) relative to the predicted 16S rRNA genes (Supplementary Fig. 4). It is intriguing that archaeal 16S rRNAs from 18 organisms showed no evidence for circularization, given that the current view is that all archaeal 16S rRNA genes go through a circular stage during processing^6^. Taking a Bathyarchaeota genome as an example, although abundant mapped transcripts extended gene ends for up to 246 bp, no transcripts supported the existence of circular 16S rRNA (Supplementary Fig. 5). Absence of circular 16S rRNAs was also observed for Thermoplasmatota and Thaumarchaeota, yet circularized 23S rRNAs in these archaea were detected (Fig. 2). Notably, all archaea in the three phyla without circular 16S rRNAs are in phylogenetic clades that are distinct from related organisms with circular 16S rRNAs (Fig. 2).

In model archaea, the first step of processing for both 16S and 23S rRNAs involves an RNA splicing endonuclease (EndA) that recognizes bulge-helix-bulge (BHB) motifs and cleaves transcribed rRNA into precursors^6,25^. Excised precursor rRNAs are then ligated by RNA splicing ligase (RtcB), generating circular intermediates^11,26^. We thus searched for EndA and RtcB in our circularized and near-complete archaeal genomes (Fig. 2), and identified them in all cases, regardless of whether or not the data supported a circularization step during 16S rRNA processing (Supplementary Figs. 6 and 7). This is not unexpected, given the observation of circular 23S rRNAs in all the archaea studied here.

We next sought EndA recognition sites, the BHB motifs, in 16S and 23S rRNAs predicted secondary structures. Consistent with the inferred RNA forms from transcript mapping, sequences with circularized 16S and/or 23S rRNAs have canonical BHB motifs in the region where circularization occurs (Supplementary Figs. 1 and 8). In contrast, sequences with non-circular 16S rRNAs do not have BHB motifs (Supplementary Figs. 5 and 9). Without a BHB motif, EndA cannot recognize and excise 16S rRNA from precursors, and as a consequence, no circular 16S rRNA transcripts would be generated.

The findings raise the question of how archaeal 16S rRNA genes without BHB motifs are processed if they cannot go through a circularization step. If the rRNA genes are arranged in a single operon and co-transcribed into 16S-23S-5S rRNA precursors, the 23S rRNAs can be excised at their BHB motifs by EndA, generating linear 16S rRNAs that may be trimmed prior to incorporation into ribosomes. However, if the rRNA genes are encoded at separate locations in genomes^27^, the 16S rRNA gene is transcribed by an independent promoter, so EndA cleavage and circularization may not be required. Although trimming is necessary to form mature molecules, it remains unclear which enzymes perform this function. Perhaps these archaea exploit RNases, analogous to bacterial and eukaryotic processing pathways^6,28,29^.

We found no correspondence between the genomic organization of the rRNA genes and whether or not their 16S rRNA genes go through a circularization step. Of 78 archaea that can generate circular 16S rRNA transcripts, 41 encode rRNA genes in operons and 35 encode separate rRNA genes. Of 18 archaea that are unable to circularize 16S rRNA transcripts, 3 have rRNA genes arranged in operons and 12 have them separated. In the other cases both for archaea that do and do not circularize their 16S rRNA transcripts, genome fragmentation precluded determination of the coding pattern.

Notably, some archaeal phyla (e.g., Bathyarchaeota) include organisms that do and do not circularize their 16S rRNA transcripts during processing (Fig. 2). We wondered whether and how this difference may relate to growth rates, as rRNA transcription is closely associated with ribosome synthesis and cell growth^30^. The GC skew profiles of circular Bathyarchaeota genomes indicated bidirectional replication from a single origin (Supplementary Fig. 10). Thus, we estimated their *in situ* replication rates using a method developed for genomes that are replicated bidirectionally^31^. Bathyarchaeota that apparently circularize their 16S rRNA transcripts had significantly higher replication rates than their counterparts (average iRep values: 1.67 vs. 1.30, *p* = 0.000015; Supplementary Fig. 11). Circular RNAs are typically harder to degrade than linear ones^32^, and the greater stability of circular rRNAs may promote ribosome synthesis and thus facilitate faster microbial growth.

Where there is evidence for an intermediate circularized form of 16S rRNA, Nanopore long-read sequencing without rRNA pre-depletion showed that the amounts of circular and linear 16S rRNA transcripts were of similar magnitudes (Fig. 3). Thus, it was surprising to note up to thousands of 23S rRNA transcripts in the circular form, with very little evidence for linear counterparts. For example, ∼6 k Nanopore transcripts showed circularization of 23S rRNA from a near-complete *Methanoperedens* genome (bMp^33^) but no transcripts indicated linear versions (Fig. 3). Given that the majority of rRNAs derive from mature ribosomes^19,30^, this finding suggests that the 23S rRNAs in circular forms in the studied archaea are within ribosomes.

**Fig. 3.**
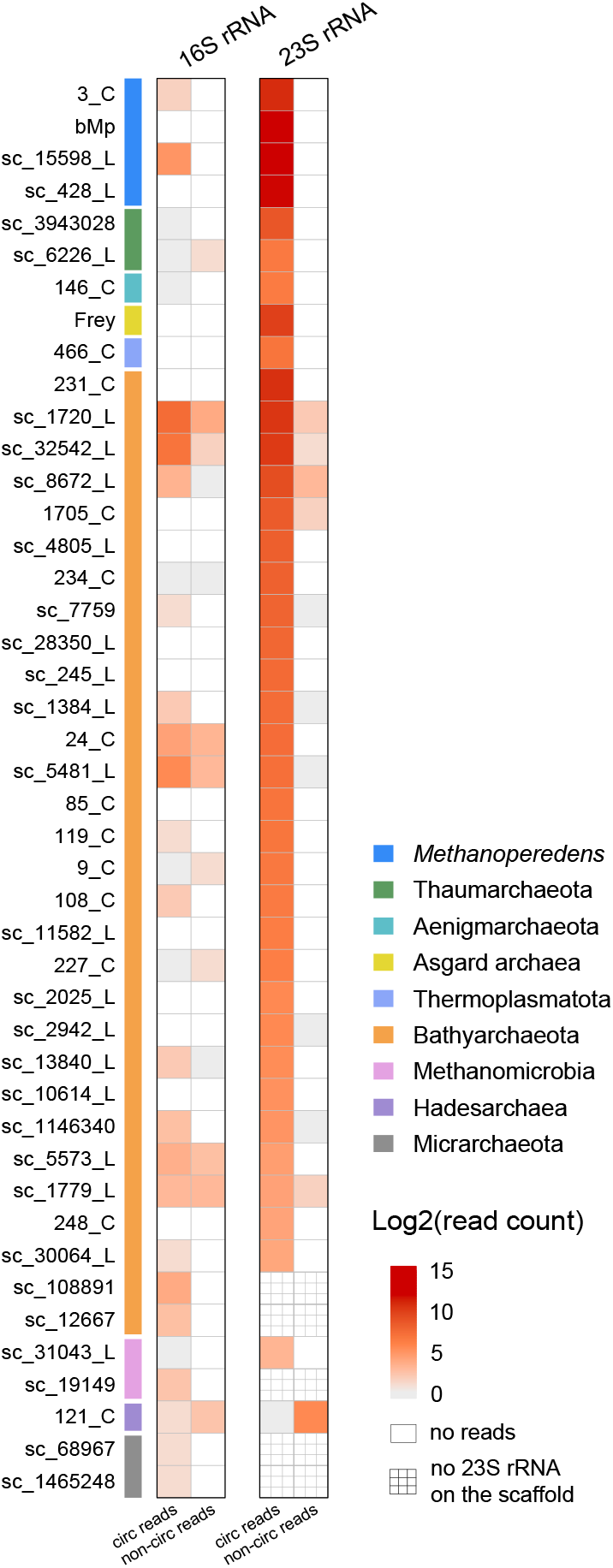
Quantitation of Nanopore transcripts indicative of different rRNA forms. Nanopore transcript reads were mapped to rRNA genes with maximum mismatches of 1%. The column “circ reads” indicates transcripts that support the circular form of rRNAs, and “non-circ reads” denotes transcripts that support the linear form (see Fig. 1). Read counts were converted logarithmically with base 2 for visualization purposes. The y-axis represents genome names and the colored bars indicate taxonomic affiliations.

To test the hypothesis that some archaeal ribosomes contain a circularized form of the 23S rRNA, we isolated large and small ribosomal subunits from the model archaeon, *Methanosarcina acetivorans*^34^, and directly sequenced the extracted 23S and 16S rRNAs. As expected based on our quantitative analysis of transcript mapping (Fig. 3), 16S rRNA in the *M. acetivorans* ribosomes is linear (Supplementary Fig. 12). However, all mapped transcript reads at the start and ends of the 23S rRNA genes (> 3k) showed discrepant read parts relative to the reference, indicative of circularization, whereas no reads extended into the flanking intergenic regions consistent with linearization (Fig. 4a). The results clearly indicate that the 23S rRNAs within the ribosomes are circular (Supplementary Fig. 13). Notably, discrepant regions start at the BHB splicing motif. The *M. acetivorans* circular 23S rRNA is 33 bp longer than the initially predicted gene and is terminated by a GNRA tetraloop that may cap the RNA sequence in circularized form (Figs. 4b and 4c).

**Fig. 4.**
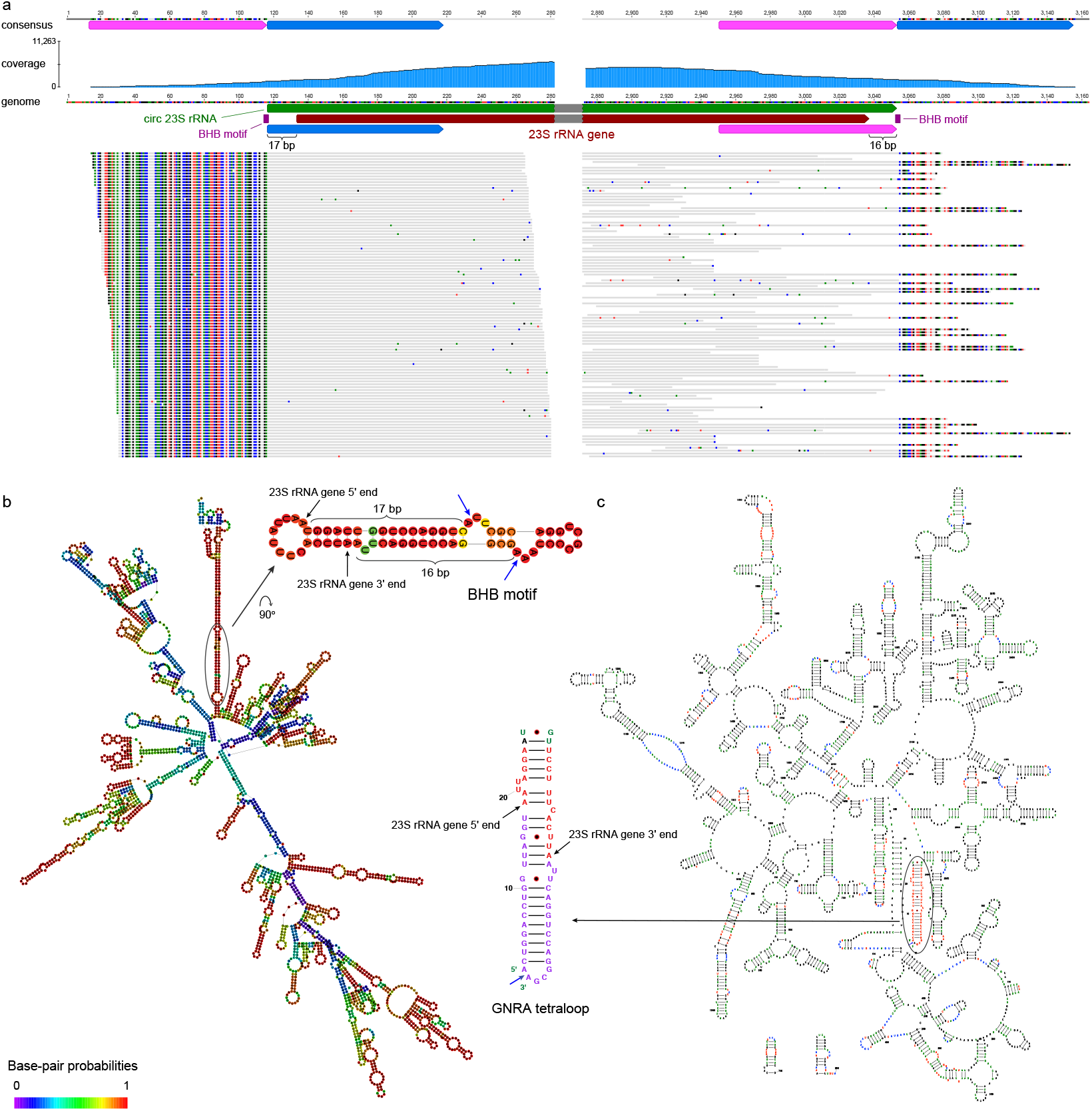
Circular 23S rRNA within the *Methanosarcina acetivorans* ribosome. (**a**) *M. acetivorans* genome mapped by 250-bp transcript reads. The dark red arrow indicates the 23S rRNA gene predicted by Rfam and the green arrow indicates its circular transcript form inferred from transcript read mapping. Purple boxes show the bulge regions in the BHB splicing motif. Pink and blue arrows demonstrate identical sequences in the genome reference and in the consensus sequence of mapped reads. Gray bars are mapped reads, in which colored dots are mismatched nucleotides to the genome. Details can be found in Supplementary Fig. 13. (**b**) Predicted secondary structure of the *M. acetivorans* 23S rRNA gene region by the Vienna RNAfold web server. Colors indicate base-pair probabilities. The ellipse region is magnified, showing the locations of the BHB motif and the 5′ and 3′ ends of the predicted 23S rRNA gene. Blue arrows indicate the cleavage sites on the BHB motif. (**c**) 2D structure of the circular 23S rRNA in the *M. acetivorans* ribosome predicted by RNAcentral and curated manually. The sequence is capped with the GNRA tetraloop located outside the 5′ and 3′ ends of the predicted 23S rRNA gene. The blue arrow indicates the site where circularization occurs. The purple nucleotides show the extra 33 bp of the circular 23S rRNA relative to the predicted gene as depicted in (**a**).

It would be interesting to verify the prediction of circularized 23S rRNAs within mature ribosomes of the uncultivated archaea studied here. To do this would require isolation of ribosomes from low-abundance archaea from extremely microbially complex wetland soil samples. We attempted to do this using several strategies (see Methods), but the efforts were unsuccessful.

To examine the potential consequences of incorporating circular 23S rRNAs into ribosomes, we analyzed the available Cryo-EM structure of a 70S ribosome from a model archaeon, *Thermococcus kodakarensis* (PDB: 6SKF)^35^. The structure shows that the 5’ and 3’ ends of the 23S rRNA form a helix and are in close proximity (Supplementary Fig. 14). However, there are several nucleotides at the ends of the 23S rRNA that are unresolved in the reported structure due to dynamics, which preclude the visualization of the circularized 5’ and 3’ ends. In contrast, the start and end of the 16S rRNA are spatially separated in the small ribosomal subunit (Supplementary Fig. 14). As circularization of 16S rRNA would disrupt the ribosome structure, it is unsurprising that we found no evidence for the presence of circularized 16S rRNA within mature ribosomes.

In combination, the extremely low incidence of linear forms of 23S RNAs and the findings for *M. acetivorans* suggest that many archaea incorporate circular 23S RNAs into their ribosomes. We considered other explanations for the lack of detectable (or appreciable) linear 23S rRNAs in cases without experimental evidence. One possibility may be the dominance of 23S rRNA pools by circular molecules not yet incorporated into ribosomes. This might be possible if the cells were growing extremely rapidly. However, Illumina and Nanopore sequencing with and without rRNA pre-depletion showed very low average transcript coverages in genomic regions that flank rRNA genes (Supplementary Fig. 15), indicating that the archaea were not growing rapidly. It would be surprising if microorganisms transcribe and accumulate, but do not use rRNA, as ribosome assembly is more rapid than rRNA transcription^30^. In conclusion, our findings indicate that circular 23S rRNAs exist within many archaeal ribosomes. This contrasts with the long-standing view that, across all the domains of life, only linear rRNAs can be assembled into ribosomes^36^.

It is interesting that some (and maybe most) archaea adopt a circularization step during rRNA processing, which does not occur in bacteria or eukaryotes. In the current study, we observed evidence for circular 23S rRNA in the ribosomes of Asgard archaea (Fig. 2), which are argued to share a common ancestor with eukaryotes^37–39^. This suggests that eukaryotic RNA processing mechanisms either evolved after divergence from Asgard archaea or were acquired from Bacteria. We also found some lineages of archaea appear not to generate a circular intermediate of 16S rRNA, establishing that the 16S rRNA processing pathway is not universal across the archaeal domain. Finally, we showed for one archaeal isolate, and inferred for many uncultivated archaea, that the ribosomes incorporate a circularized form of the 23S rRNA. This is a revision of our understanding of ribosome biology and raises the intriguing question of how the inclusion of circular molecules may be advantageous. One conceivable benefit is the greater stability and durability of ribosomes, as increased interactions between the 5’ and 3’ ends of 23S rRNA can improve the thermostability of the large subunit^40^. It may also confer resistance to exonucleases and thus reduce ribosome turnover for lower metabolic costs of organisms^41^.

How archaea assemble circular 23S rRNAs into ribosomes remains unclear. One possibility could be that ribosomal proteins bind to their corresponding sites on pre-circularized 23S rRNAs directly and proceed with the assembly in the 5′-3′ direction^42–44^. However, circular RNAs impose stronger topological constraints relative to linear counterparts that limit their conformational flexibility^45^, which may impede ribosome biogenesis. The other possibility is that linear precursors of circular 23S rRNAs interact with ribosomal proteins and assemble into the large ribosomal subunit, and their free ends further ligate to each other on mature ribosomes. Future imaging experiments based on pure cultures may address these possibilities.

## Methods and Materials

### Nucleic acid extraction and sequencing

Samples were collected at various depths from wetland soil in Lake County, California, USA. Approximately 5.0 g of soil samples at 75 cm and 140 cm depths were used for DNA extraction using Qiagen DNeasy PowerMax Soil Kit. This DNA was sequenced using PacBio HiFi long reads to generate new metagenomic datasets for recovery of high-quality rRNA genes and for comparison with previous short-read datasets. Other samples from the same site at depths from 40 cm to 115 cm were used for RNA extraction by Qiagen RNeasy PowerSoil Total RNA Kit. A subset of RNA samples from 50 cm, 90 cm, 100 cm, and 115 cm were sequenced using Nanopore long reads^33^, and others from 40 cm, 60 cm, 80 cm, and 100 cm were sequenced using Illumina PE150 short reads^23^. Nanopore RNA sequencing did not pre-deplete rRNAs. In brief, all the RNA were polyadenylated using poly(A) polymerase (NEB, M0276L) and purified by GeneJET RNA Cleanup and Concentration Micro Kit (ThermoFisher, K0841). The poly(A)-tailed RNA was reverse transcribed and amplified to generate full-length cDNA according to the Oxford Nanopore Technologies PCR cDNA Synthesis (PCS109) protocol. The cDNA amplicons were barcoded (EXP-NBD114), pooled and sequenced using FLO-PRO114M flow cells on PromethION.

### Processing of metagenomic and metatranscriptomic sequencing data

All raw sequencing reads were processed to filter out low-quality data using BBDuk (https://jgi.doe.gov/data-and-tools/software-tools/bbtools/). PacBio HiFi reads were assembled using hifiasm-meta (v0.13-r308)^46^. Scaffolds ≥ 1 kb were binned by a combination of CONCOCT (v1.1.0)^47^, MaxBin2 (v2.2.7)^48^, MetaBAT2 (v2.15)^49^, and VAMB (v3.0.2)^50^. Open reading frames (ORFs) and ribosomal RNA genes in assembled scaffolds were predicted using Prodigal (v2.6.3)^51^ and Rfam 14^52^, respectively.

### Identification of genomes that can circularize rRNAs

A total of 11,710 16S rRNA genes were identified from the wetland soil site (Supplementary Fig. 3). We removed genes that were shorter than 1,400 bp, likely fragmental genes, and that were longer than 2,000 bp, likely false positives, which resulted in 4,580 putative complete 16S rRNA genes. These genes were clustered at ≥ 99% identity and ≥ 90% coverage, yielding 2,576 species-level 16S rRNA genes. As Nanopore RNA sequencing did not involve the pre-depletion of rRNAs, we mapped processed Nanopore transcript reads to the species-level 16S rRNA genes using Minimap2 (v2.28-r1209) with default parameters in the “map-ont” mode^53^. Resulting SAM files were converted to BAM formats using SAMtools (v1.17)^54^, followed by calculations of transcript coverages and covered fractions using CoverM (v0.6.1)^55^. We further discarded genes that have coverages < 5 and/or covered fractions < 50%, leading to a final collection of 1,125 species-level 16S rRNA genes. We retrieved the corresponding genomes of these genes and mapped both Nanopore and Illumina transcript reads to them. Mapping files were visualized in Geneious Prime 2024.0.4 (https://www.geneious.com) to figure out the forms of rRNAs, as depicted in Fig. 1. We also preliminarily classified the 16S rRNA genes using the SILVA ACT web service^56^.

### Gene annotation and phylogeny construction

Gene analyses were conducted on self-circular PacBio genomes and near-complete genomes that include most single-copy genes and < 20 scaffolds. RNA splicing endonuclease (EndA) was identified with HMMER (v3.3) using profile hidden Markov Models (HMM) PF01974 and PF02778^57^. RNA ligase (RtcB) was identified according to HMM PF01139.

Proteins of interest were blasted against the Genbank nr database to recruit homologues^58^. Query sequences and references were aligned using MAFFT (v7.453)^59^. Alignments were trimmed by trimAl (v1.4.rev15) and then used for phylogeny construction by IQ-TREE (v1.6.12) with automatically-selected best-fit models^60,61^. Trees were visualized and decorated on the iTOL server^62^.

Predicted 16S rRNA genes were blasted against the SILVA database (v138)^63^ and the GTDB SSU database (v214)^64^ for the recruitment of homologous sequences. Two *Escherichia coli* 16S rRNA genes (A14565.1 and AB045730.1) and three *Bacillus subtilis* genes (AB016721.1, AB042061.1 and AB055007.1) were used as the outgroup sequences. Further alignment, trimming, and phylogeny construction were the same as described above.

### Measurement of *in situ* replication rates and theoretical minimal doubling time

Metagenomic sequencing reads from samples at depths of 60-175 cm were mapped to self-circular and near-complete Bathyarchaeota genomes using BBMap with a minimal read identity of 97%. Generated SAM files were sorted by SAMtools (v1.17)^54^ and input into iRep (v1.10) to calculate *in situ* genome replication rates and profile GC skew^31^. Theoretical minimal doubling times of these genomes were estimated using the gRodon package with the “metagenome_v2” mode^65^.

### Transcript quantification of rRNAs and flanking genomes

To calculate the transcript abundances of circular and linear rRNA forms, Nanopore transcript reads were mapped to rRNA genes with maximum mismatches of 1%. Reads extending beyond rRNA gene ends were analyzed, according to Fig. 1, to quantify circular versus linear forms. Read counts were converted logarithmically with base 2 for visualization purposes.

To calculate the transcript abundances of genomes flanking rRNA genes, rRNA gene sequences were masked with ‘N’, and the modified genomes were mapped with Nanopore and Illumina transcript reads using maximum mismatches of 3%. Average transcript coverages were calculated by dividing total mapped transcript bases by flanking genome length. The resulting values were converted logarithmically with base 2. To avoid biases caused by too short reference length, only genomes and/or scaffolds ≥ 50 kb were included for analysis.

### Modeling of RNA 2D structures

Secondary structures of rRNA sequences were predicted on the RNAfold web server^66^. To locate the possible BHB splicing motifs, an extension of sequences flanking the rRNA genes (e.g., 200 bp) was included. rRNA 2D structural diagrams with consistent, reproducible, and recognizable layouts were modeled with RNAcentral^67^ and manually curated using XRNA-React^68^.

### Extraction and sequencing of RNA from *Methanosarcina acetivorans* ribosomes

*M. acetivorans* ribosomes were isolated from the pure cultures as described previously^34^. RNA was extracted from the ribosomes using Qiagen RNeasy Mini Kit (Cat. No. 74104). Libraries were prepared and sequenced using the Illumina MiSeq 250PE platform in the QB3 Genomics, UC Berkeley. Metatranscriptomic reads were first processed to filter out low-quality data using BBDuk and then mapped to the *M. acetivorans* 16S and 23S rRNA genes, which are predicted by Rfam 14^52^, using BBMap with a minimal read identity of 97%. Matched reads were re-mapped to the *M. acetivorans* genome (GenBank: AE010299.1) with a loose identity of ≥ 70% in Geneious Prime 2024.0.4 (https://www.geneious.com) to visualize transcript reads that support circular or linear rRNA forms (Fig. 1). Secondary structures of rRNAs were predicted as described above.

### Attempt for ribosome isolation from complex soil samples

Approximately 5.0 g of soil samples were resuspended in 25 mL ribosome buffer A (20 mM Tris-HCl pH 7.5, 100 mM NH^4^Cl, and 10 mM MgCl^2^) and sonicated in a Q500 sonicator (Qsonica) for 20 minutes. Another portion of samples were resuspended in 12 mL PowerBead solution and 0.5 mL Solution SR1 with PowerMax beads from the Qiagen RNeasy PowerSoil Total RNA Kit and vortexed at maximum speed for 30 minutes. The solutions were clarified by centrifugation and loaded onto sucrose cushions containing 24 mL buffer B (20 mM Tris-HCl pH 7.5, 500 mM NH^4^Cl, and 10 mM MgCl^2^) with 0.5 M sucrose and 17 mL buffer C (20 mM Tris-HCl pH 7.5, 60 mM NH^4^Cl, and 6 mM MgCl^2^) with 0.7 M sucrose. Ribosomes were pelleted at 57,000 x *g* for 16 hours at 4°C in a Ti-45 rotor (Beckman-Coulter). Pellets were resuspended in ribosome dissociation buffer (20 mM Tris-HCl pH 7.5, 60 mM NH^4^Cl, and 1 mM MgCl^2^) and were loaded onto a 25-40% (w/v) sucrose gradient in ribosome dissociation buffer. The gradients were centrifuged in a SW-32 rotor (Beckman-Coulter) at 97,000 x *g* for 16 hours at 4°C. Gradients were analyzed on a Biocomp piston fractionator.

## Data availability

Nanopore transcript data can be accessed in the NCBI database under BioProject accession: PRJNA1119519 (BioSample: SAMN41664889-SAMN41664893)^33^. A subset of Illumina transcript data has been deposited in the NCBI database under BioProject accession: PRJNA1050611 (BioSample: SAMN41840554-SAMN41840557)^23^. Other Illumina metagenomic and metatranscriptomic raw sequences, along with PacBio long reads data, have been deposited under the BioProject accession of PRJNA1247515. The genomes can be accessed at: https://ggkbase.berkeley.edu/circ_rRNA_paper_related.

## Acknowledgments

This work was supported by the Chan Zuckerberg Climate Fund and by the funding from the U.S. Department of Energy (DOE) Chemical Sciences, Geosciences, and Biosciences Division under Contract DE-AC02-05CH11231. Ribosome isolation experiments were supported by the NSF Center for Genetically Encoded Materials (C-GEM; CHE 2002182).

## Author Contributions

The study was designed by L-D.S. and J.F.B. L-D.S. took soil samples, extracted RNA, and processed metatranscriptomic sequencing data. L-D.S. analyzed rRNA forms, gene functions and phylogenies, transcript abundances, and genome replication patterns and rates. P.I.P. and L-D.S. performed 2D structural analysis on rRNAs. A.J.N. performed 3D structural analysis on *T. kodakarensis* ribosomes. A.J.N. isolated ribosomes from *Methanosarcina acetivorans* cell pellets with oversight by D.D.N. and J.H.D.C. L-D.S. extracted RNA from *M. acetivorans* ribosomes and analyzed transcriptomic sequencing data. R.S. processed metagenomic sequencing data and provided bioinformatic support. L-D.S. and J.F.B. wrote the paper with input from all the authors.

