## Supplementary Information for "Circular 23S rRNA within archaeal ribosomes"

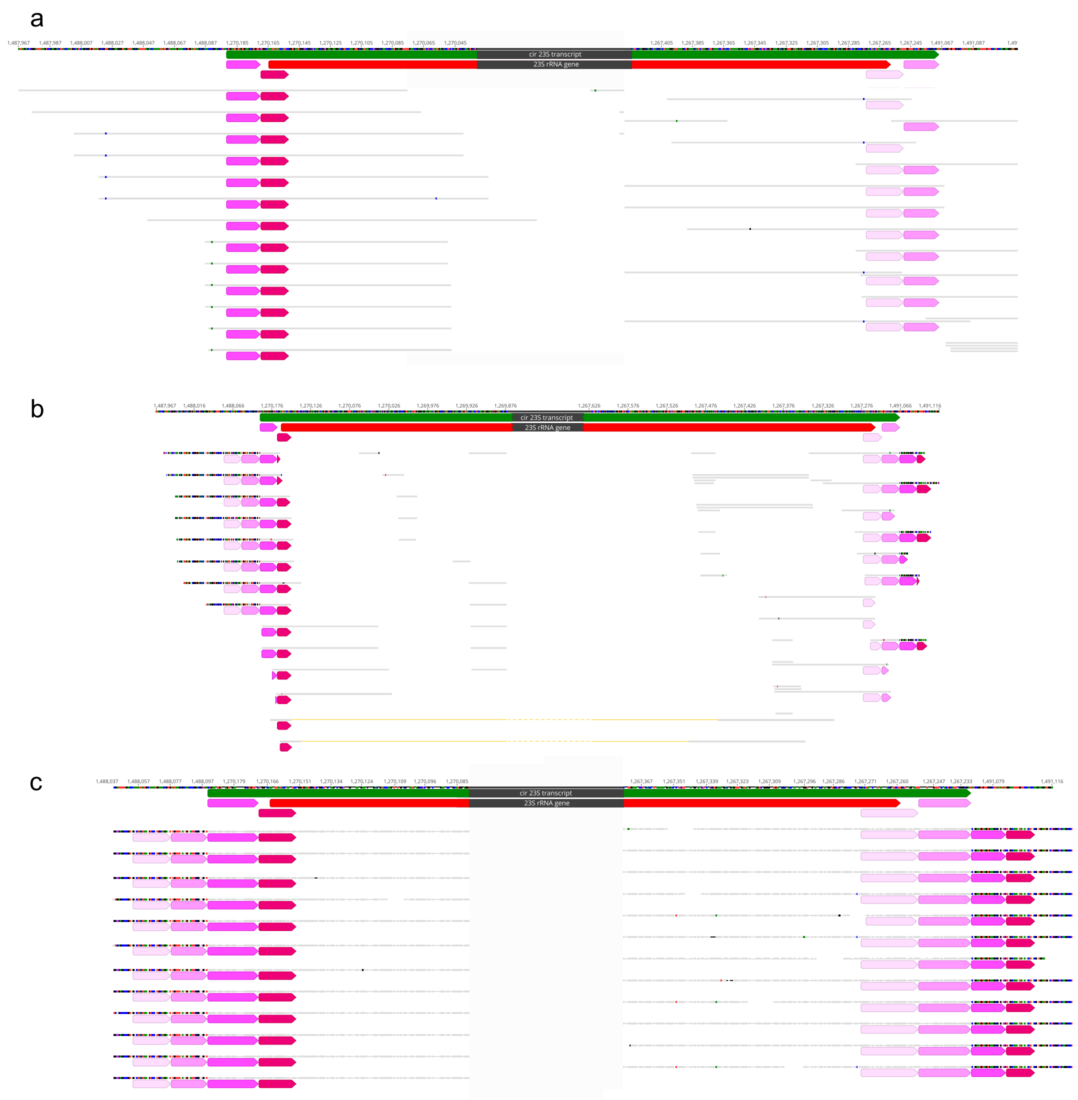


**Supplementary Fig. 1 Example of read mapping showing circular rRNA transcripts instead of rRNA genes.** This is a *Methanoperedens* genome fragment mapped by Illumina metagenomic reads (**a**), Illumina metatranscriptomic reads (**b**), and Nanopore transcripts (**c**). The long red arrow in the genome reference indicates the predicted 23S rRNA gene and the green arrow shows the region of circular transcript inferred from transcript mapping. Gray bars are mapped reads, below which colored arrows indicate sequences ≤ 1 mismatch.

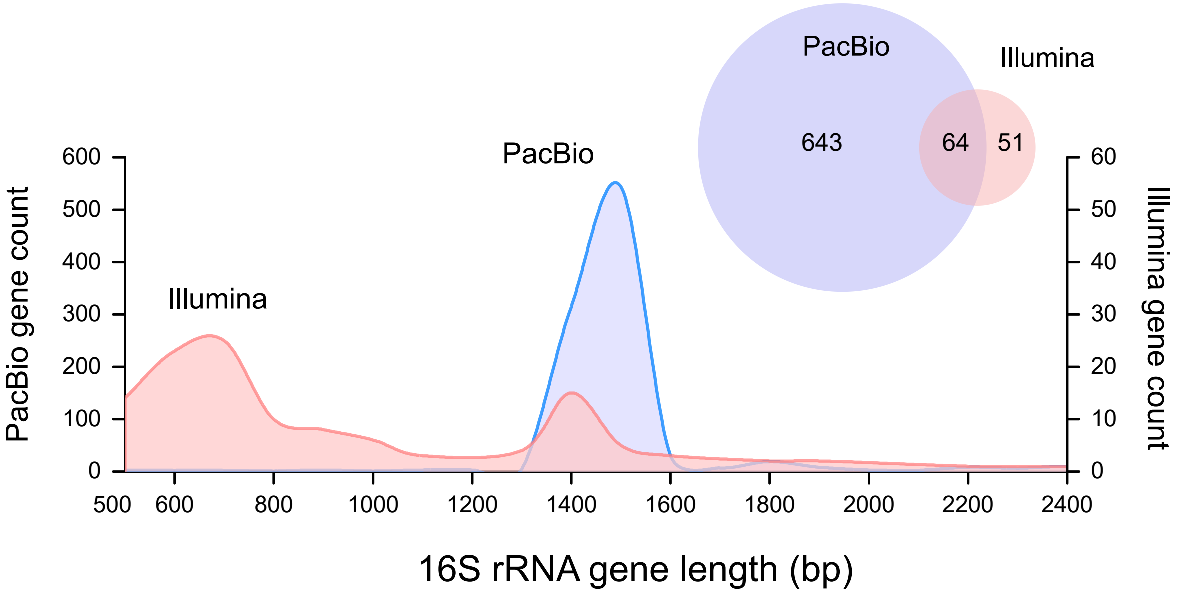


**Supplementary Fig. 2 Example of 16S rRNA genes assembled from PacBio and Illumina sequencing using the same soil sample.** The total data sizes are 38.6 Gb and 36.2 Gb, respectively, for PacBio and Illumina PE250 sequencing. The Venn diagram indicates the counts of distinct and shared 16S rRNA genes clustered at 99% of identity.

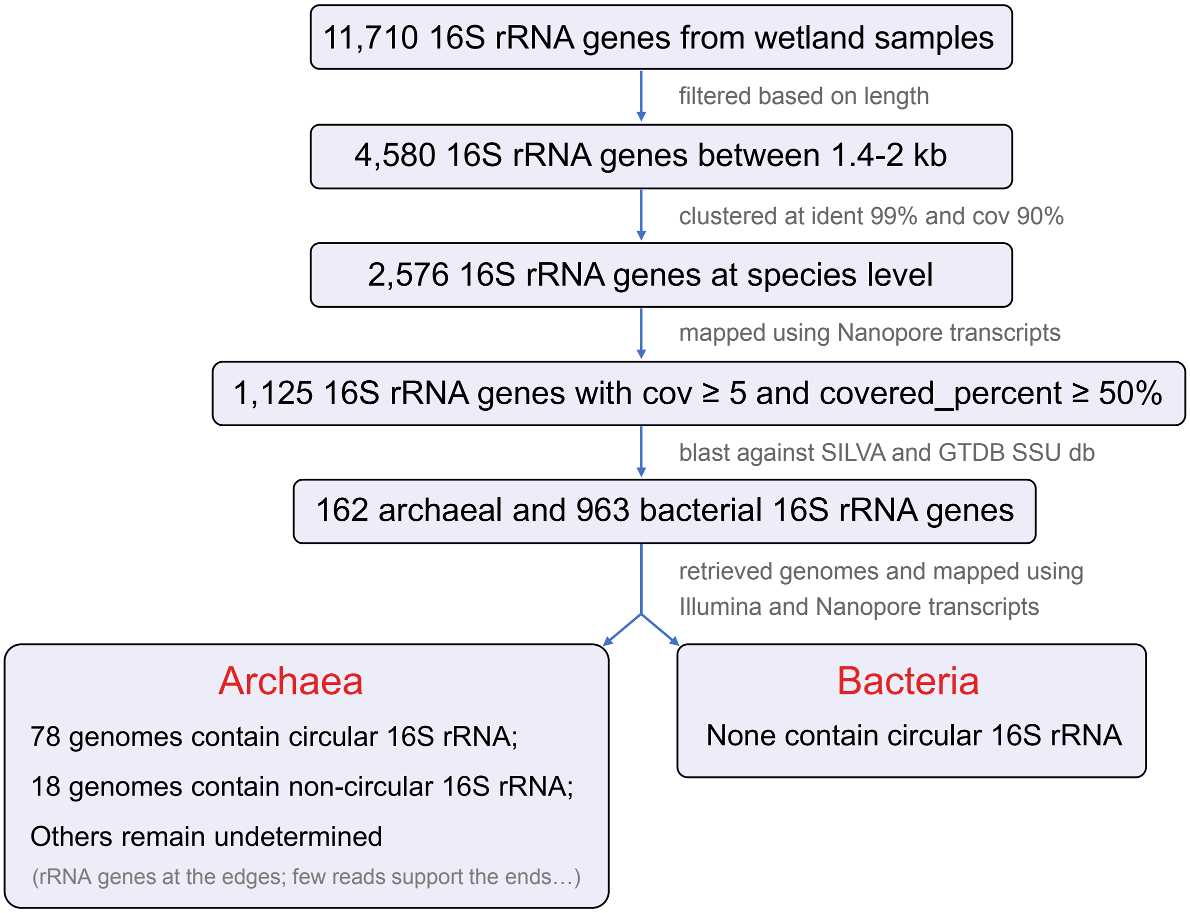


**Supplementary Fig. 3 Workflow of exploring 16S rRNA forms.**

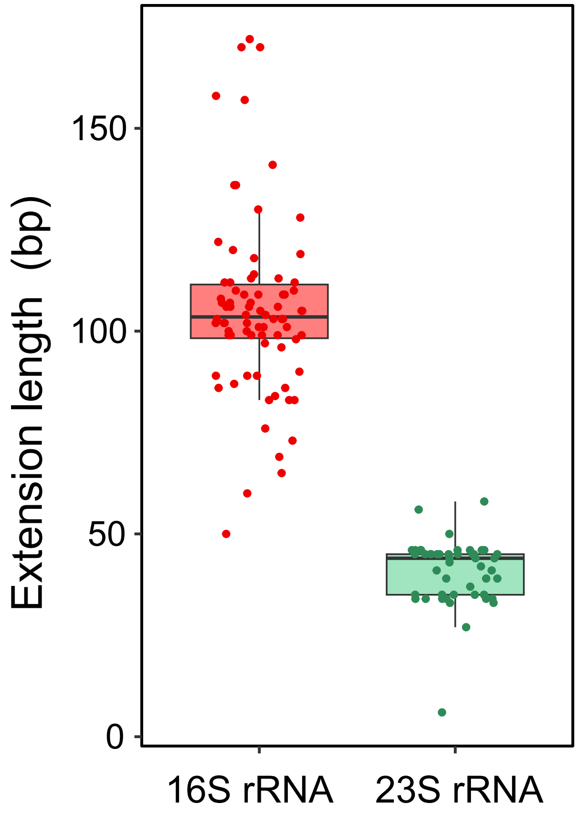


**Supplementary Fig. 4 Extension length of circular rRNA transcripts relative to genes predicted *in silico*.** Colored dots indicate circular transcripts that contain 16S (n=78) and 23S rRNAs (n=48). Box plots show lower and upper quartiles and median values in each transcript group.

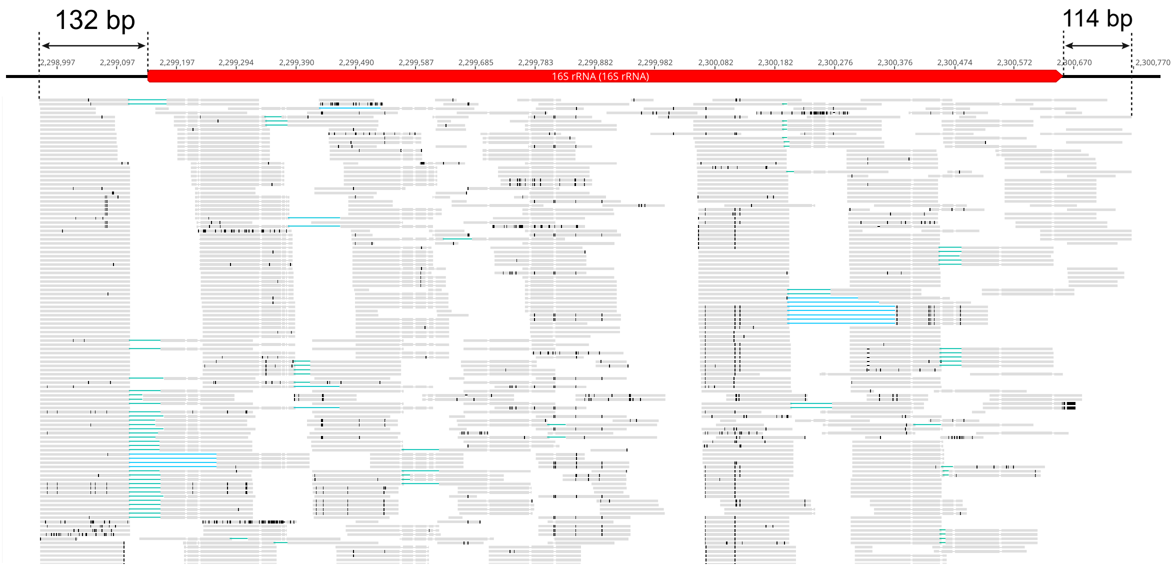


**Supplementary Fig. 5 Example of linear 16S rRNA transcript.** This is a Bathyarchaeota genome fragment mapped by Illumina transcripts with maximum mismatches of 3%. The long red arrow in the genome reference indicates the predicted 16S rRNA gene. Gray bars are Illumina transcripts matching the reference and vertical black lines indicate mismatches.

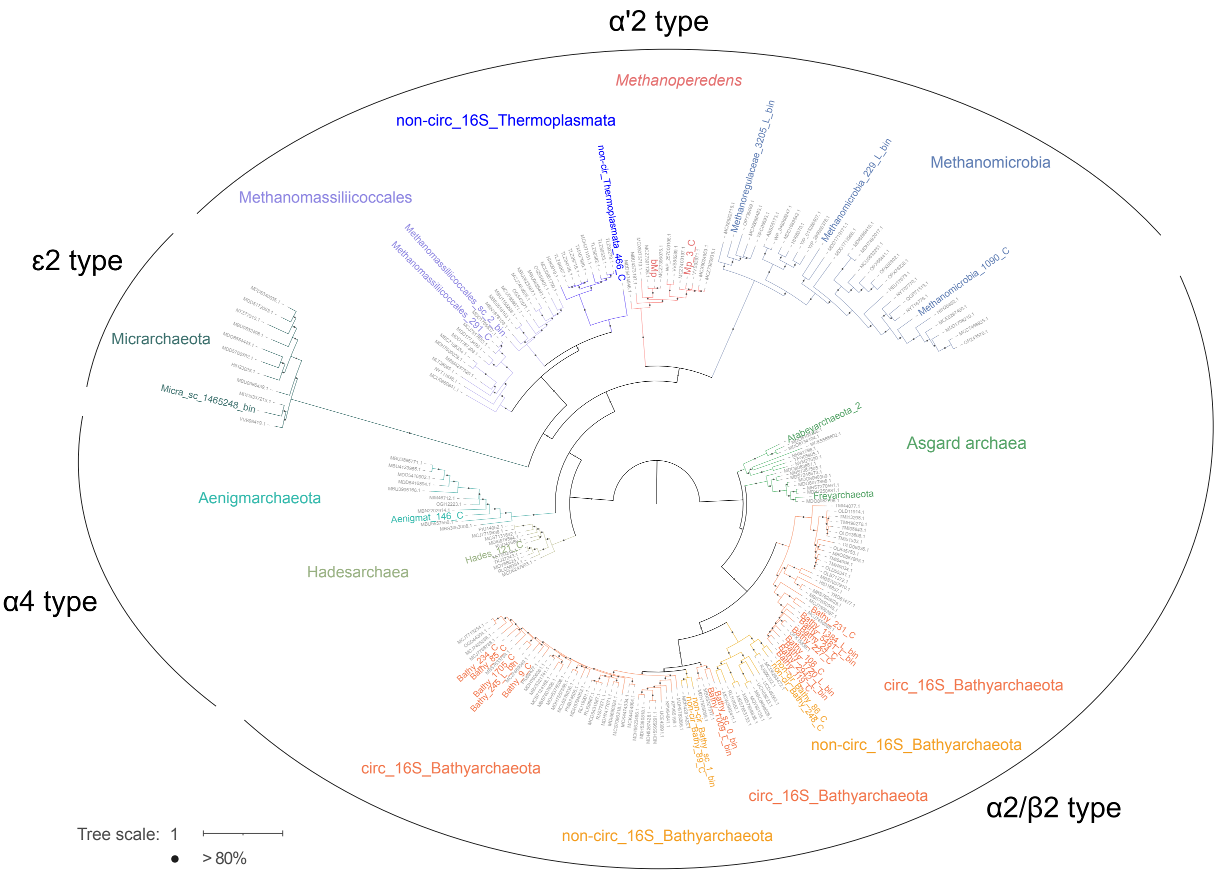


**Supplementary Fig. 6 Phylogeny of RNA splicing endonuclease (EndA).** Sequences identified in our potentially complete or near-complete genomes are colored. Genome information can be found in Fig. 2. Support values were calculated based on 1000 replicates.

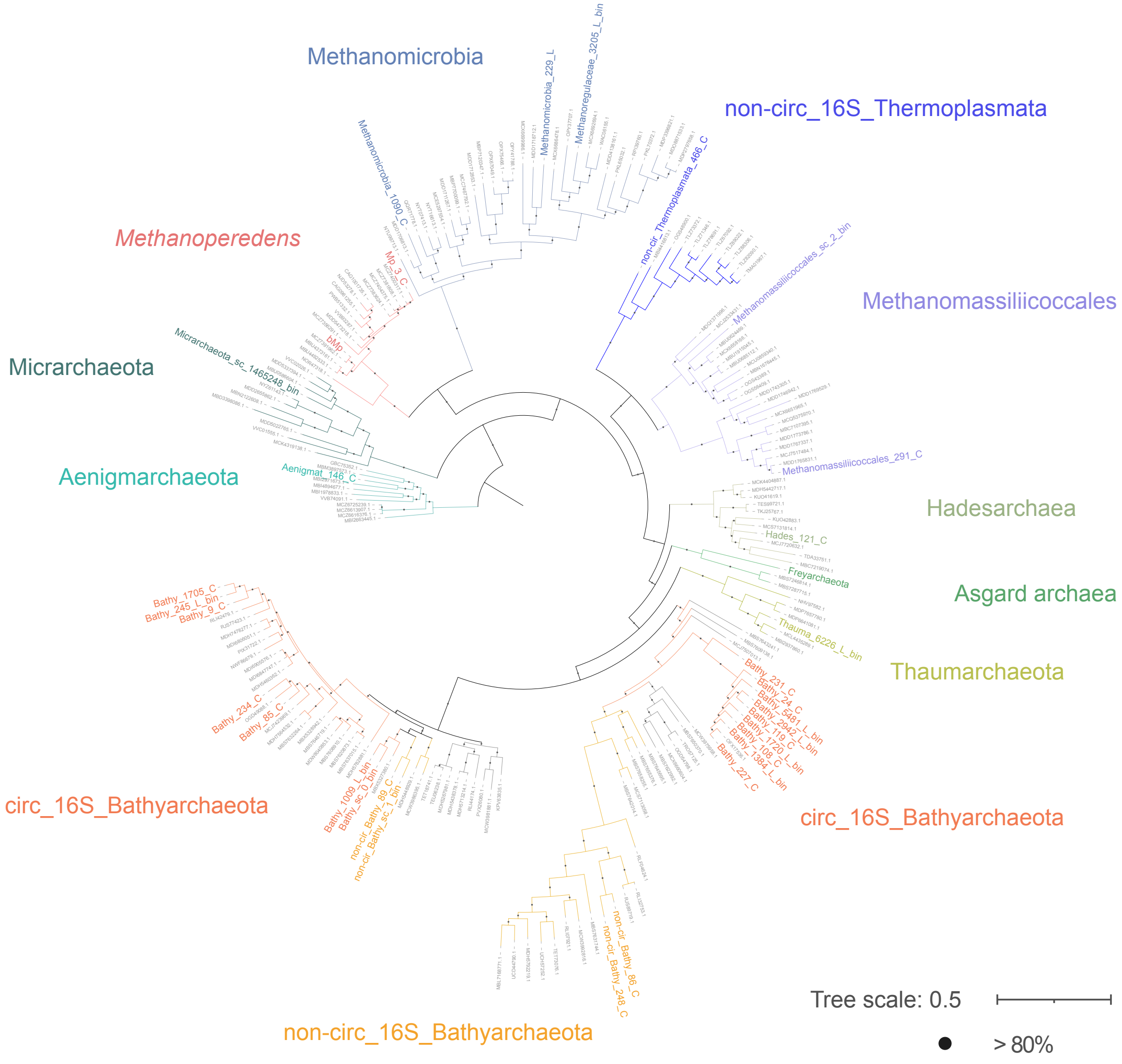


**Supplementary Fig. 7 Phylogeny of RNA ligase (RtcB).** Sequences identified in our potentially complete or near-complete genomes are colored. Genome information can be found in Fig. 2. Support values were calculated based on 1000 replicates.

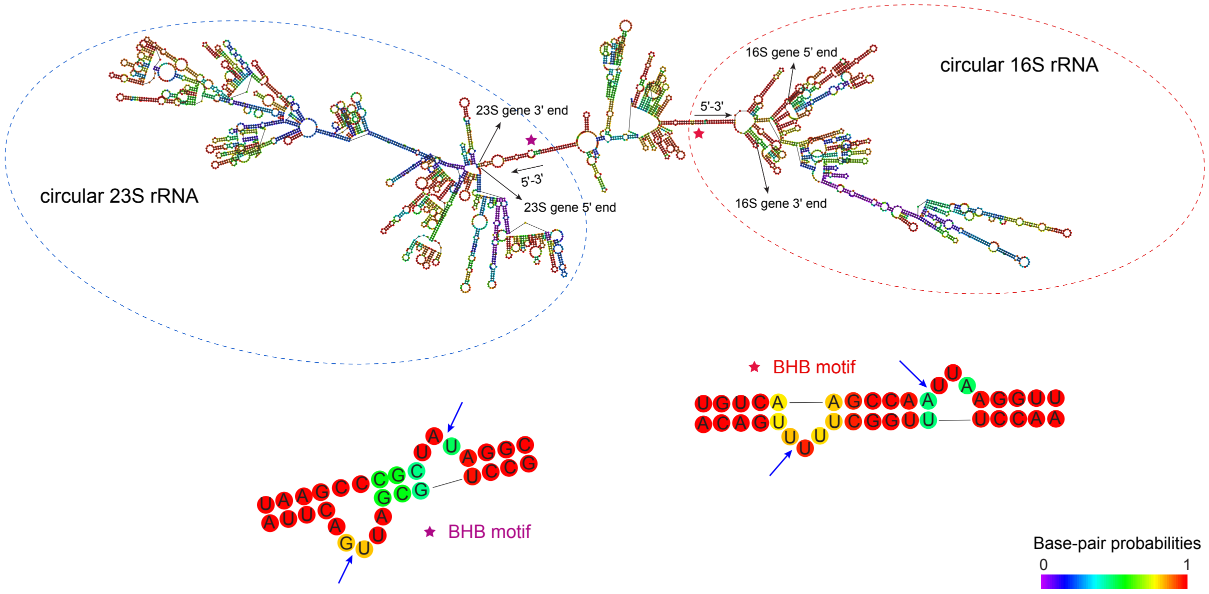


**Supplementary Fig. 8 Predicted secondary structure of the polycistronic transcript generating circular 16S and 23S rRNAs.** The sequence is from a *Methanoperedens* genome as illustrated in Supplementary Fig. 1. BHB splicing motifs in the transcript are labeled with stars and detailed at the bottom. Blue arrows indicate the cleavage sites on the BHB motifs.

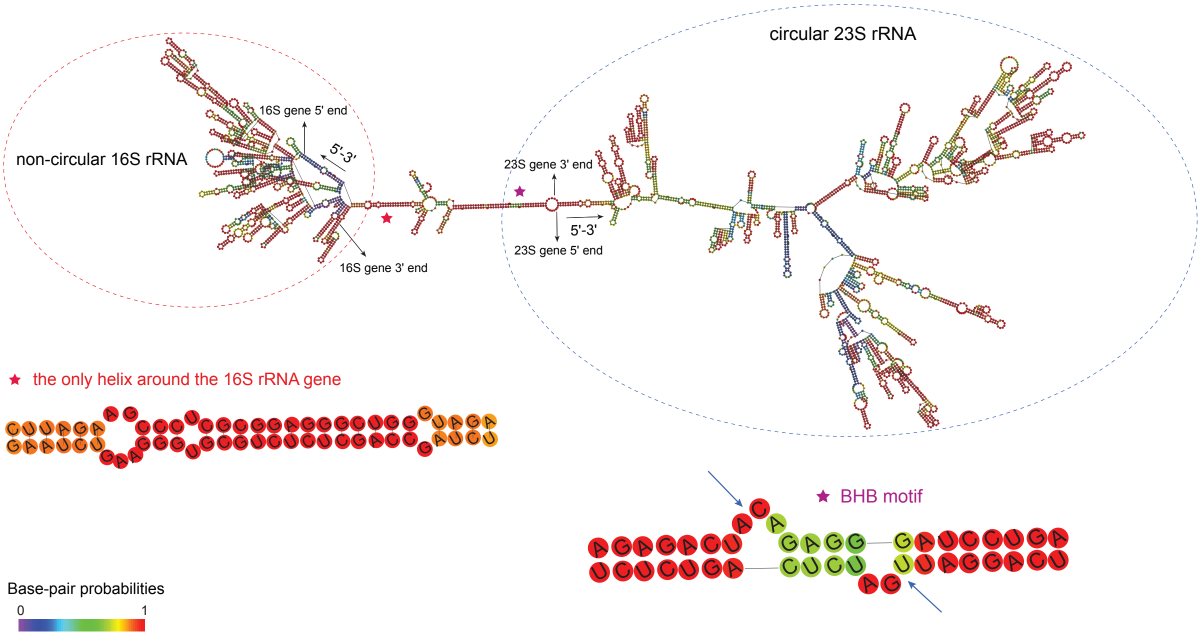


**Supplementary Fig. 9 Predicted secondary structure of the polycistronic transcript generating non-circular 16S and circular 23S rRNAs.** The sequence is from a Bathyarchaeota genome as illustrated in Supplementary Fig. 5. The only helix around the 16S rRNA gene and the BHB motif around the 23S rRNA gene are labeled with stars and detailed at the bottom. The blue arrow indicates the cleavage sites on the BHB motif.

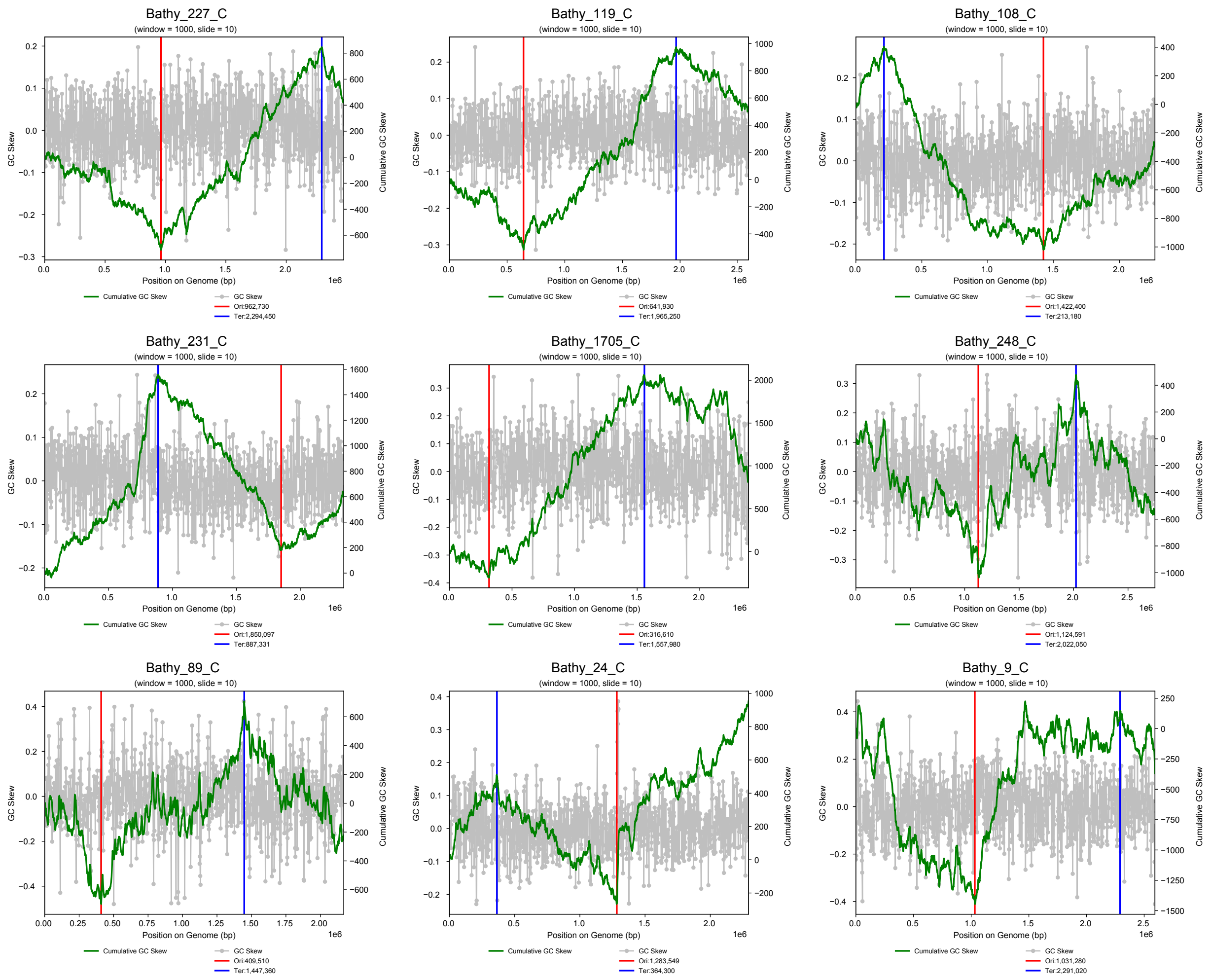


**Supplementary Fig. 10 GC skew profiles of circular Bathyarchaeota genomes.** Gray dots and green lines indicate GC skew and cumulative GC skew across genomes. Red and blue lines show the inferred locations of replication origins and termini.

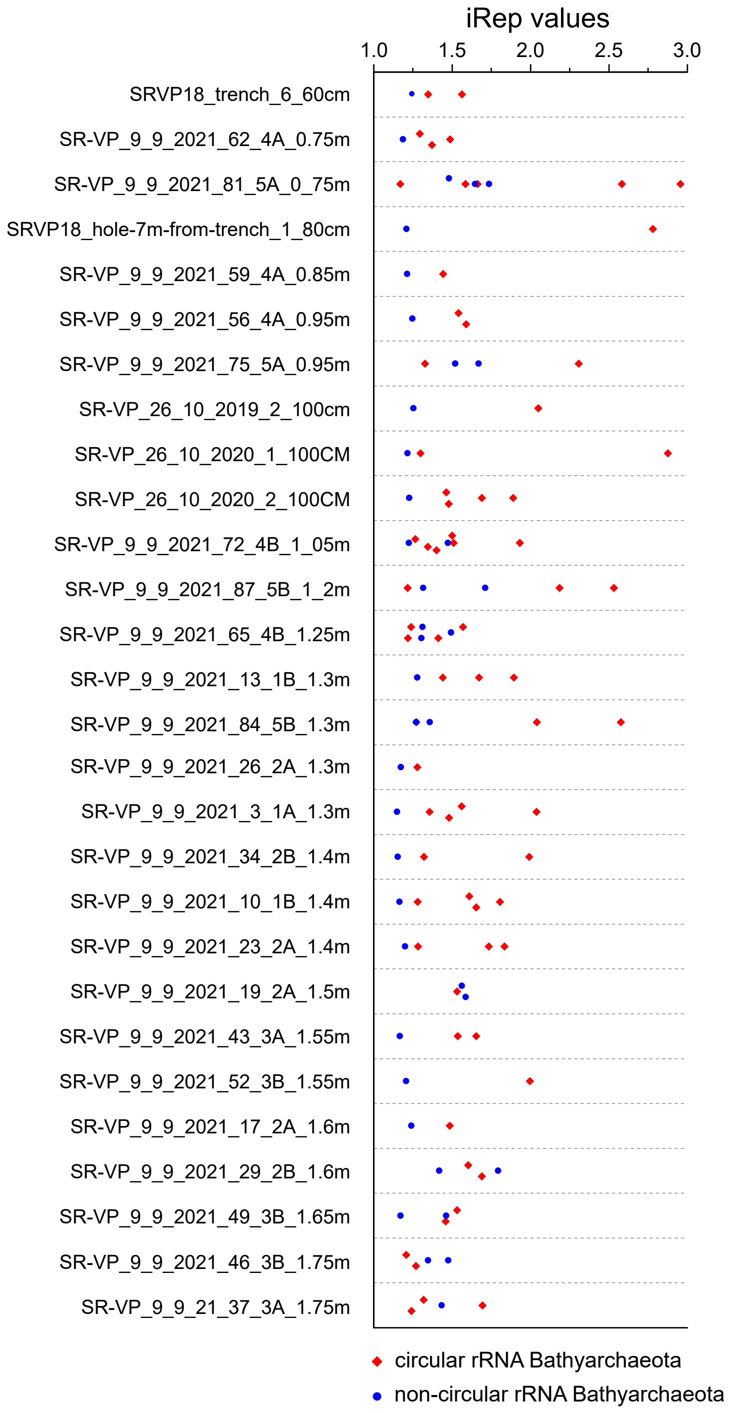


**Supplementary Fig. 11 *in situ* replication rates of Bathyarchaeota genomes generating circular or non-circular 16S rRNA intermediates.** Samples are shown only if both genome types co-existed there. Significant differences in iRep values are observed between the two genome types ($p$=0.000015), using the Wilcoxon signed-rank test.

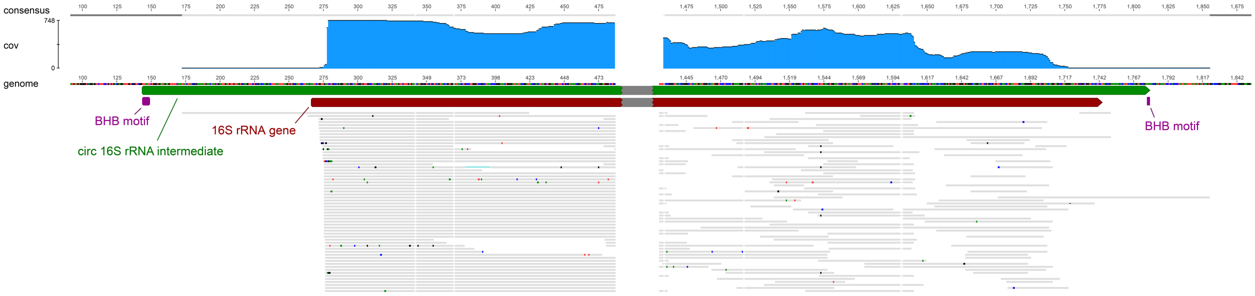


**Supplementary Fig. 12 Linear 16S rRNA within the *Methanosarcina acetivorans* ribosome.** *M. acetivorans* genome was mapped by 250-bp transcript reads. The dark red arrow indicates the 16S rRNA gene predicted by Rfam, and the green arrow indicates its circular transcript intermediate inferred from transcript read mapping. Purple boxes show the bulge regions in the BHB splicing motif. Gray bars are mapped reads, in which colored dots are mismatched nucleotides to the genome. No reads support the circularization of 16S rRNA within the *M. acetivorans* ribosomes.

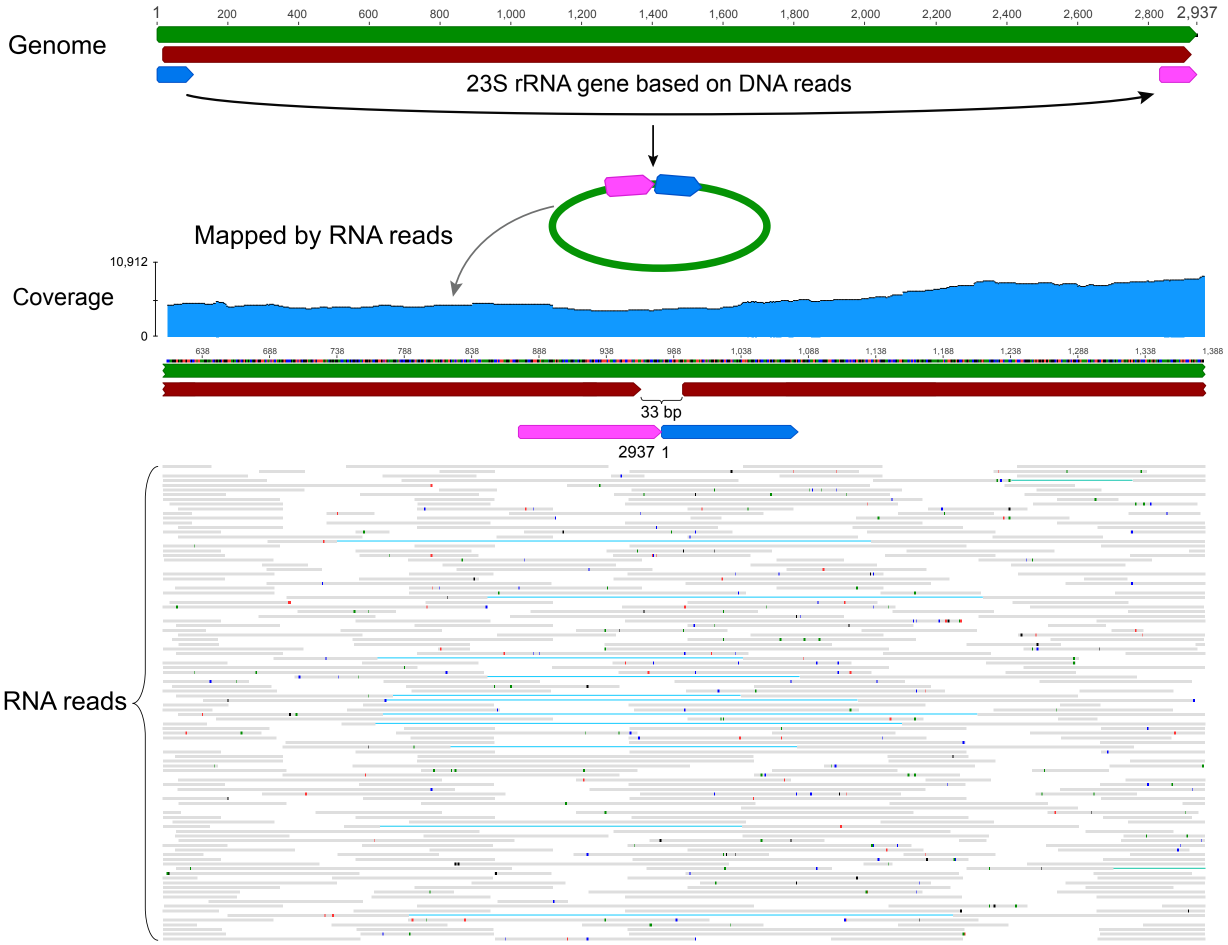


**Supplementary Fig. 13 Transcript mapping of circular 23S rRNA within the *Methanosarcina acetivorans* ribosome.** The blue and pink arrows correspond to those in Fig. 4a. The region labeled with arrows in the permuted circular 23S rRNA is mapped with RNA reads. Gray bars show mapped transcript reads wherein colored dots are mismatched nucleotides to the reference.

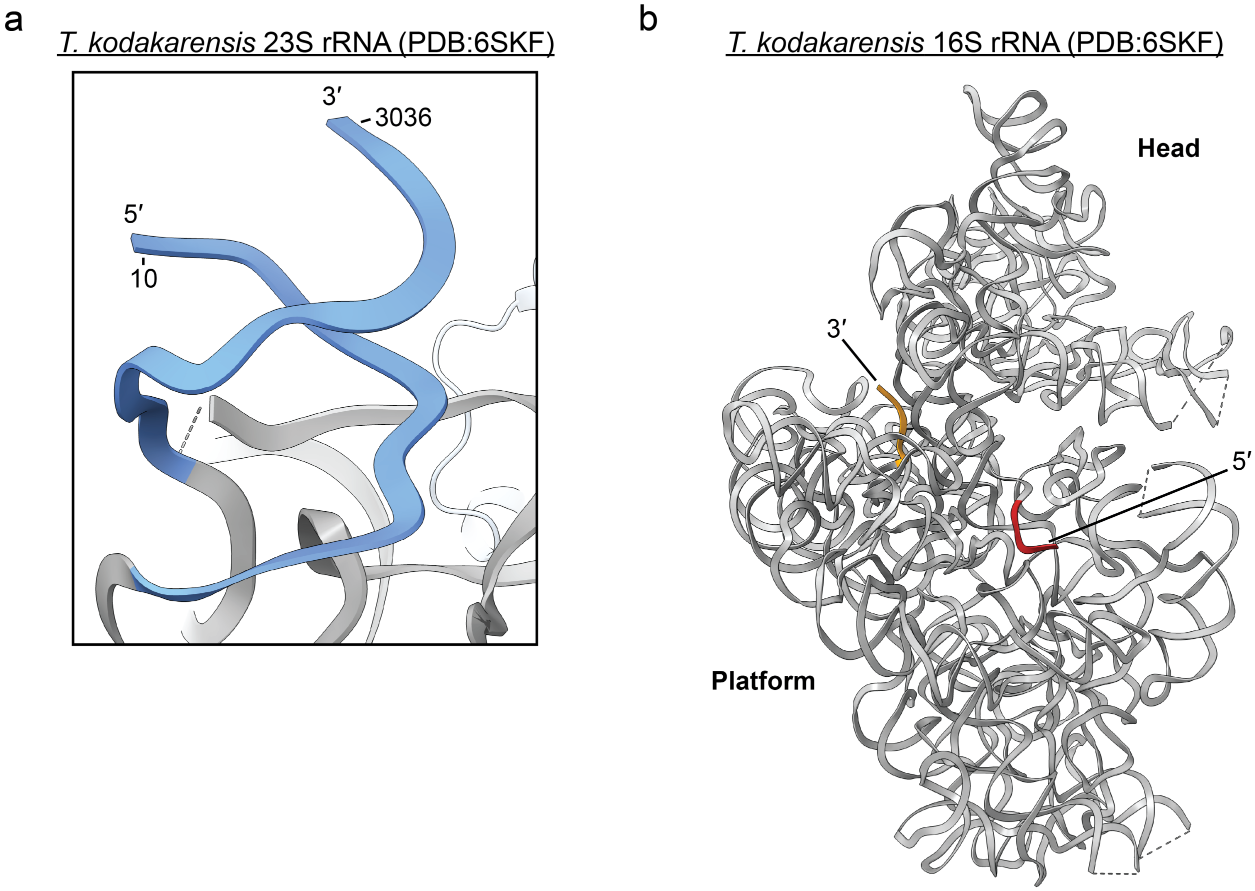


**Supplementary Fig. 14 Structural analysis of the *Thermococcus kodakarensis* ribosome.** (**a**) The 5′ and 3′ ends of the *T. kodakarensis* 23S rRNA are shown in blue. The residue numbers where the 23S rRNA model begins and ends in PDB 6SKF are indicated. (**b**) 5′ (red) and 3′ (orange) ends are highlighted in the 16S rRNA from PDB 6SKF.

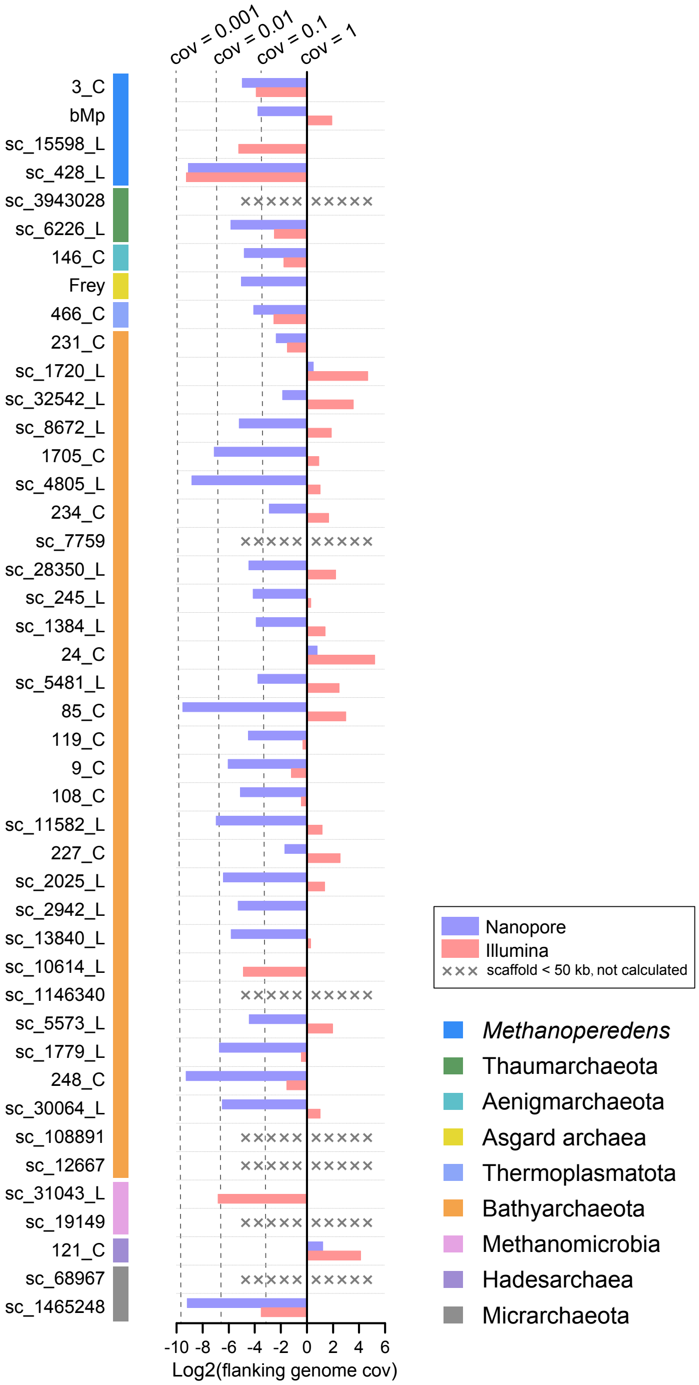


**Supplementary Fig. 15 Average transcript coverages of genomes flanking rRNA genes.** Both Nanopore and Illumina transcripts were mapped to flanking genomes with maximum mismatches of 3%. Average coverages were calculated by dividing total mapped transcript bases by flanking genome length and converted logarithmically with base 2. To avoid biases caused by too few genes, only genomes (scaffolds) ≥ 50 kb were included.
